# Abnormally increased intrinsic neural timescales in sensory and default mode networks in cocaine use disorder

**DOI:** 10.1101/2025.10.22.683885

**Authors:** Xiaoling Tang, Tianqi Di, Jintao Sheng, Yan Sun, Wenbo Luo, Mingming Zhang

## Abstract

Cocaine use disorder (CUD) is associated with abnormal structural and functional brain changes. However, the neurodynamics and molecular underpinnings remain unclear. In this study, we mapped whole-brain intrinsic neural timescales (INTs), reflecting temporal neural processing, using resting-state functional magnetic resonance imaging data from 44 CUD patients and 44 healthy controls (HC). CUD showed increased INTs in visual, somatomotor, and default mode networks compared with HC. Mediation analysis linked local INTs abnormalities to altered dorsal attention network neurodynamics, associated with inhibitory control deficits. Notably, these changes were primarily correlated with alterations in gamma-aminobutyric acid type A receptors and the noradrenaline transporter. Machine learning classifiers based on INTs achieved a maximum accuracy of 75.5% in distinguishing CUD from HC, with a generalization accuracy of 65.0% on an independent dataset. This study elucidates aberrant neural mechanisms underlying CUD and highlights INTs as promising diagnostic biomarkers for clinical detection and intervention.

**Teaser:** Spatiotemporal neuroscience reveals intrinsic neural timescale disruptions underlying cocaine use disorder, offering novel diagnostic biomarkers.

## Introduction

Cocaine use disorder (CUD), marked by compulsive drug-seeking driven by alterations in the reward system (*1*), represents a growing global health concern, with more than 25 million people using cocaine worldwide in 2023 (*2*). Cocaine use significantly impairs cognitive functions, including attention, inhibition, and emotional regulation (*3*–*5*), and these deficits are closely linked to structural and functional brain changes. For example, individuals with CUD exhibit reduced cortical thickness in the bilateral insula, prefrontal cortex, and posterior cingulate cortex, alongside increased thickness in the bilateral temporal pole (*6*). Cocaine cue exposure engages multiple emotion regulation networks, including the salience and executive control systems, with altered activation patterns linked to craving and impaired inhibitory control (*7*). Resting-state functional magnetic resonance imaging (rs-fMRI) provides a means to examine aberrant neural activity in CUD (*8, 9*). Zhao et al. (*10*) reported increased connectivity between the visual network (VN) and dorsal attention network (DAN), as well as reduced connectivity between the default mode network (DMN) and limbic network (LN). However, most existing studies have primarily focused on structural changes or static connectivity, with limited exploration of the spatiotemporal dynamics of brain activity in CUD.

Brain networks can rapidly reorganize within seconds to minutes, reflecting shifts in mental states and internal regulation. (*11, 12*). These rapid changes are crucial for understanding and diagnosing psychiatric disorders. (*13, 14*). Importantly, global dynamics arise from the activity of local regions (*15*). However, most research has focused on large-scale functional networks, paying less attention to the temporal properties of local dynamics. This gap limits our understanding of how network reconfiguration contributes to neuropsychiatric conditions. Examining the temporal fluctuations of local brain regions may offer new insights into the neural mechanisms underlying CUD (*16*).

Intrinsic neural timescales (INTs) offer a temporal framework for analyzing brain information processing and reveal abnormal neural dynamics in psychiatric disorders, thereby contributing to understanding their pathophysiology. Unimodal regions, such as the primary visual cortex, generally exhibit shorter INTs that support rapid perception and response processing. In contrast, transmodal regions, including the central executive network and DMN, display longer INTs that facilitate integration and higher-order processing (*17*). This unimodal-to-transmodal hierarchy reflects the functional organization of the brain (*18, 19*) and is essential for integrating and interpreting external information (*20*). At rest, INTs demonstrate a stable topological structure, but they reorganize dynamically during task performance (*21*). Such adaptive temporal integration supports higher cognitive functions, including visual perception, attention, and working memory (*22*). INTs show high reliability and are increasingly applied in psychiatric research (*23, 24*).

Disruptions in the INT hierarchy have been reported across psychiatric disorders. In autism spectrum disorder, INTs decrease in sensory and visual cortices but increase in the right caudate compared with healthy controls (HC) (*25*). Patients with schizophrenia show shorter INTs in primary sensory and motor regions relative to HC (*26, 27*). In male tobacco use disorder, significant reductions in INTs have been identified in the control network, DMN, and sensorimotor areas compared with HC (*28*). Shorter INTs in emotion-related brain regions, including the bilateral insula, left amygdala, and left temporal pole, have been observed in individuals with internet gaming disorder compared with HC (*29*). Although INTs are increasingly recognized as biomarkers for mental disorders, their alterations in CUD and their potential diagnostic value remain unclear. Distinct INT patterns have also been reported across subtypes of obsessive-compulsive disorder (*30*), and substance use such as cocaine and methamphetamine can elicit opposing neural activity patterns (*31*). These findings suggest that different addiction disorders may exhibit unique neurodynamic abnormalities. The present aimed to characterize the distribution of INTs in CUD, providing a basis for future cross-addiction comparisons and clarifying disorder-specific neurobiological mechanisms.

Recent studies have validated INTs as potential neuroimaging biomarkers for psychiatric disorders. Machine learning models using INTs have achieved 95.5% accuracy in classifying disorders of consciousness (*32*), 81.4% for major depression, and 76% for epilepsy (*33, 34*). These findings highlight INTs’ potential in diagnosing and classifying brain disorders. However, local intrinsic neural dynamics in CUD remain poorly understood, and whether INTs can reliably distinguish individuals with CUD from HC has yet to be established. Developing robust and generalizable neuroimaging biomarkers may help clarify the neurobiology and pathophysiology of CUD.

Brain neural dynamics are closely coupled with neurotransmitter distribution (*35*). Neurotransmitters shape the time scale of information processing by modulating the balance of excitation and inhibition within neural networks (*36*). Several potential relationships between INTs and neurotransmitter pathways have been identified, which may be crucial for understanding psychiatric disorders. Exploring the link between INTs and neurotransmitters may bridge macro-level neural dynamics with micro-level molecular mechanisms, offering insights into the pathology of mental disorder. In patients with depression, reduced INTs have been associated with monoamine neurotransmitters, including serotonin, noradrenaline, and dopamine (*37*). In tobacco addicts, abnormal INTs have been linked to dopaminergic, acetylcholine, and μ-opioid receptor systems (*28*).

Ricard et al. (*38*) combined functional connectivity data with dopamine D2/3 receptor positron emission tomography and demonstrated that patients with CUD exhibited widespread DMN-subcortical connectivity reductions that strongly correlated with dopamine receptor density distribution. Nevertheless, the relationship between neurotransmitter distribution and INT patterns in CUD remains unclear.

The present study investigated abnormal changes in INTs in CUD at both voxel-wise and network levels and examined their associations with clinical data (Figure 1). To explore the biomolecular basis of these alterations, spatial correspondences between INT changes and multiple neurotransmitter receptor distributions were assessed. Finally, rs-fMRI was combined with machine learning to evaluate the utility of INTs as a neural marker for distinguishing individuals with CUD from HC.

**Fig. 1.**
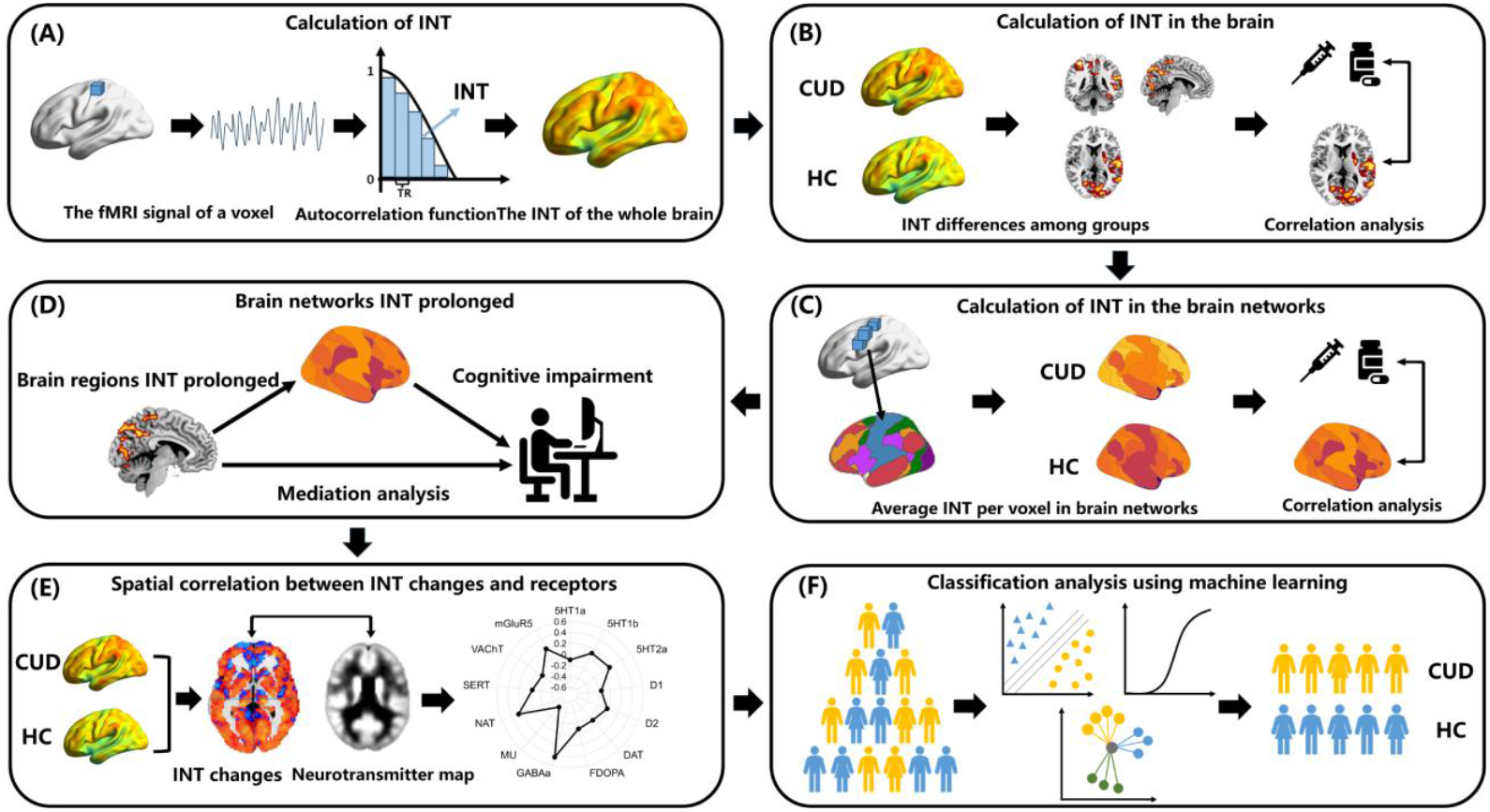
The protocol for data analysis. **(A)** fMRI signals were extracted from each voxel, the INTs were calculated using the autocorrelation function, and the whole-brain INTs distribution is subsequently visualized. **(B)** Whole-brain analysis was performed to compare the differences in INTs between CUD and HC in local brain regions. **(C)** The INTs of brain networks was calculated by averaging INTs values across voxels, and the results were visualized. **(D)** A mediation model was used to analyze the relationships between local brain region INTs, brain network INTs, and cognitive impairments in CUD. **(E)** The JuSpace toolbox was used to analyze the spatial correlations between INTs changes and neurotransmitters. **(F)** Machine learning was used to distinguish between CUD and HC. fMRI: functional magnetic resonance imaging. INTs: intrinsic neural timescales. CUD: cocaine use disorder. HC: healthy controls.

## Results

### Demographic and clinical characteristics

The demographic and clinical characteristics of the CUD and HC groups are summarized in Table 1. No significant differences in age (*t*_(88)_ = 0.33, *p* = 0.74, Cohen’s d = 0.07), gender (*χ*^2^ _(1)_ = 0.08, *p* = 0.78, Phi = 0.03), education (*t*_(88)_ = -0.40, *p* = 0.69, Cohen’s d = - 0.09). These findings indicate that the two groups were well-matched on key demographic variables, minimizing potential confounding effects in subsequent analyses.

**Table 1.** Demographic and clinical characteristics of both groups. CUD: cocaine use disorder. HC: healthy controls. FD: framewise displacements. CCQN: cocaine craving questionnaire now. NA: not applicable.

| Variable | CUD | HC | t or $\chi^2$ | *p* | Cohen's d or Phi |
| --- | --- | --- | --- | --- | --- |
| Age (years) | 32.33 ± 5.98 | 31.82 ± 8.36 | 0.33 | 0.74 | 0.07 |
| Gender (male/female) | 38/7 | 37/8 | 0.08 | 0.78 | 0.03 |
| Education (years) | 11.67 ± 2.90 | 11.93 ± 3.32 | -0.40 | 0.69 | -0.09 |
| Mean FD (mm) | 0.13 ± 0.06 | 0.13 ± 0.06 | < 0.001 | 0.99 | < 0.001 |
| Duration of cocaine use (years) | 11.09 ± 6.74 | NA | NA | NA | NA |
| Age of first cocaine use (years) | 21.13 ± 5.73 | NA | NA | NA | NA |
| Time since last use (days) | 12.43 ± 12.19 | NA | NA | NA | NA |
| CCQN | 141.23 ± 44.76 | NA | NA | NA | NA |

### Whole-brain mapping reveals prolonged INTs in CUD

To examine the effects of CUD on hierarchical temporal integration and regional variation, whole-brain INT spatial maps were compared between patients with CUD and HC. Both groups displayed the typical unimodal-to-transmodal INT hierarchy (Figure 2A): INTs were longer in the frontal and parietal cortices and shorter in the auditory, visual, and sensorimotor regions. This hierarchical structure was consistently observed in analyses without global signal regression (GSR; Figure S1). Whole-brain analysis revealed that, compared with the HC group, the CUD group showed prolonged INTs in the left middle temporal gyrus, right superior parietal gyrus, left fusiform gyrus, left pallidum, and left cerebellum_6 (*p* ≤ 0.05, Threshold-Free Cluster Enhancement (TFCE)-false discovery rate (FDR) corrected; Table 2 and Figure 2B). No regions showed reduced INTs in CUD relative HC. At the level of functional modules, the significant clusters with increased INTs in CUD primarily overlapped with the ventral attention network (VAN) (Dice coefficient (DC) = 0.115), somatomotor network (SMN) (DC = 0.116), DAN (DC = 0.079) and DMN (DC = 0.079; Figure 2B). In whole-brain analyses without GSR, peak coordinates shifted, but the main affected regions remained consistent (Figure S2 and Table S1).

**Table 2.** Significant group differences in INTs. AAL: Anatomical Automatic Labeling. MNI = Montreal Neurological Institute. CUD: cocaine use disorder. HC: healthy controls. INTs: intrinsic neural timescales. L: left; R: right.

| Region | AAL | Peak MNI Coordinate |  |  | Voxels | t value |
| --- | --- | --- | --- | --- | --- | --- |
|  |  | X | Y | Z |  |  |
| Temporal_Mid_L | 85 | -51 | 3 | -30 | 830 | 5.06 |
| Parietal_Sup_R | 60 | 33 | -42 | 60 | 809 | 5.00 |
| Fusiform_L | 55 | -30 | -51 | -15 | 716 | 5.05 |
| Pallidum_L | 75 | -15 | 6 | 3 | 20 | 3.64 |
| Cerebellum_6_L | 99 | -21 | -57 | 27 | 14 | 3.46 |

**Fig. 2.**
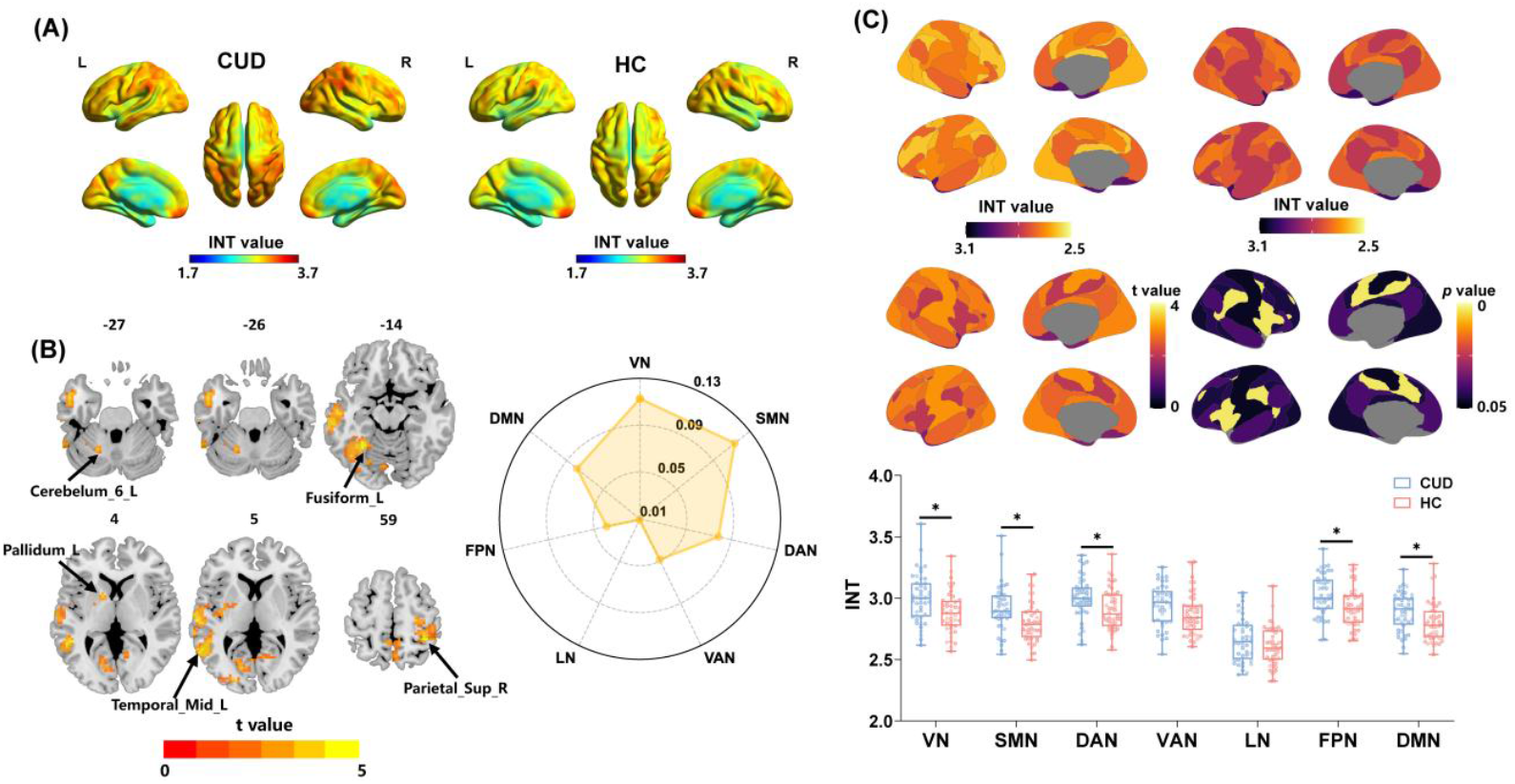
The distribution of INTs and differences between CUD and HC groups. **(A)** The spatial distribution of INTs in CUD and HC Groups. **(B)** The brain regions with significant differences in INTs between the CUD and HC groups. The statistically significant threshold was set at *p* < 0.05 (two-tailed), TFCE-FDR corrected. **(C)** The distribution pattern of the brain network INTs in the CUD and HC groups. CUD: cocaine use disorder. HC: healthy controls. INTs: intrinsic neural timescales. L: left; R: right. VN: visual network. SMN: somatomotor network. DAN: dorsal attention network. VAN: ventral attention network. LN: limbic network. FPN: frontoparietal control network. DMN: default mode network. * *p* < 0.05, FDR corrected.

### Abnormally prolonged INTs in sensory and DMN in CUD

To directly assess the involvement of functional modules in CUD, average INTs were extracted from each functional network and compared between the CUD and HC groups. INTs were significantly longer in the CUD group with in the VN (*t*_(86)_ = 2.94, *p* = 0.013, after FDR corrected, Cohen’s d = 0.63), SMN (*t*_(86)_ = 3.05, *p* = 0.013, after FDR corrected, Cohen’s d = 0.65), DAN (*t*_(86)_ = 2.75, *p* = 0.013, after FDR corrected, Cohen’s d = 0.59), frontoparietal control network (FPN) (*t*_(86)_ = 2.85, *p* = 0.013, after FDR corrected, Cohen’s d = 0.61), and DMN (*t*_(86)_ = 2.60, *p* = 0.015, after FDR corrected, Cohen’s d = 0.55; Figure 2C). No significant differences were observed in the VAN (*t*_(86)_ = 2.02, *p* = 0.055, after FDR corrected, Cohen’s d = 0.43) and LN (*t*_(86)_ = 1.43, *p* = 0.156, after FDR corrected, Cohen’s d = 0.31). In analyses without GSR, significant differences in the VN and SMN were replicated (Figure S3). A difference in the DMN was also detected, although it did not survive FDR correction.

Taken together with the whole-brain analysis, these findings suggest that abnormal INT prolongation in CUD is primarily localized to the VN, SMN, DAN, and DMN relative to HC.

### Abnormal INTs in the pallidum and DMN persist during abstinence in CUD

To examine the relationship between altered INTs and clinical features in CUD, average INT values were extracted from regions of interest (ROIs) that showed significant differences between the CUD and HC groups. Correlation analysis revealed that INT values in the left pallidum were positively associated with the duration of abstinence (*r* = 0.369, *p* = 0.048, after FDR corrected; Figure 3A). INT values in the DMN also showed a positive correlation with abstinence duration (*r* = 0.362, *p* = 0.017); although this association did not survive FDR correction (corrected *p* = 0.054). Collectively, these findings suggest that the neural impairments associated with cocaine use are not transient but may exhibit long-term stability.

**Fig. 3.**
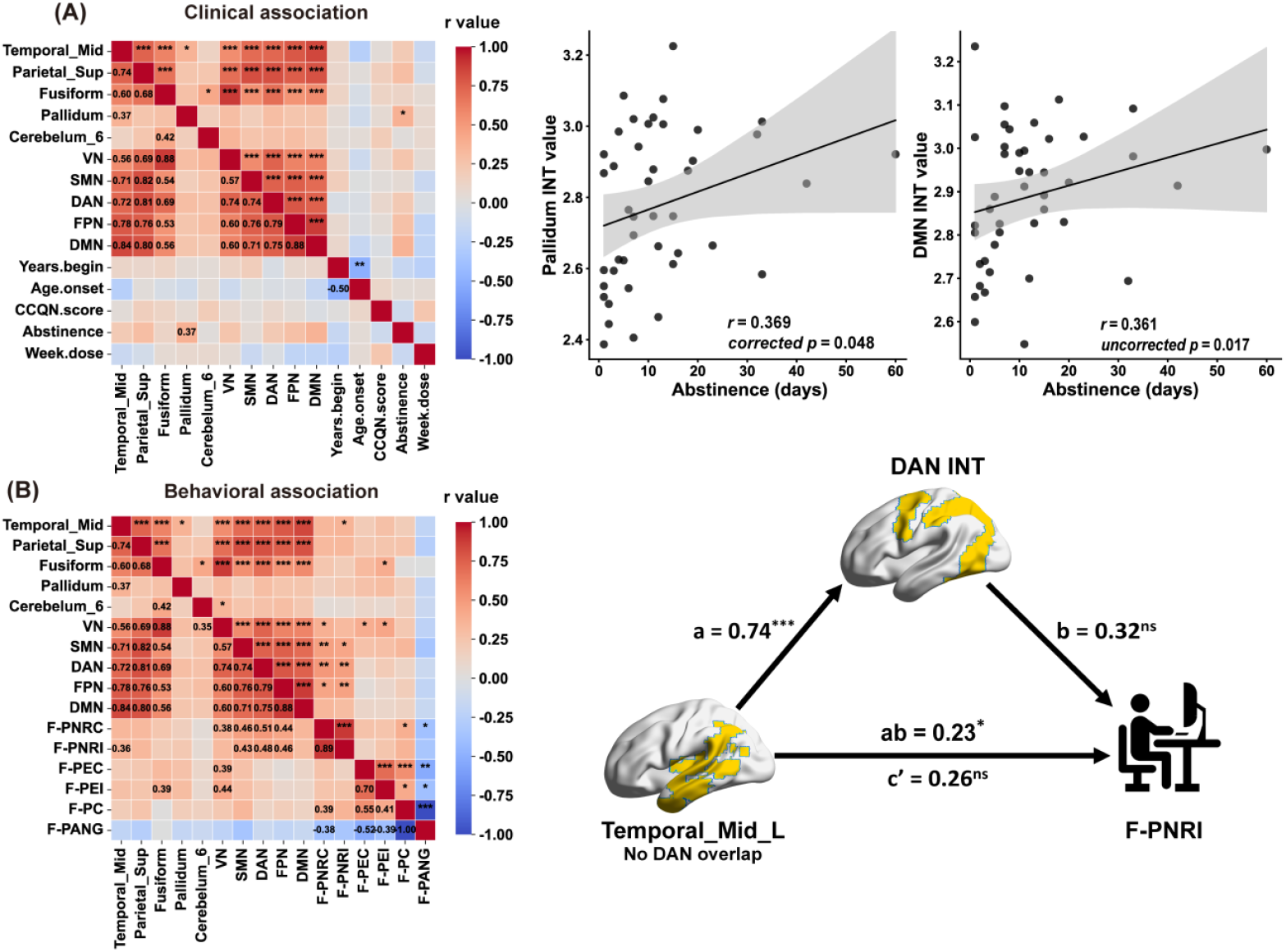
The relationship between INTs and clinical and behavioral information in CUD. **(A)** Correlation matrix of INTs in brain regions/networks with clinical information. The INTs in the left pallidum and the DMN showed a significant positive correlation with abstinence. **(B)** Correlation matrix of INTs in CUD with behavior information. A mediation model with regional INTs as the independent variable, DAN INTs as the mediator, and behavioral information as the dependent variable. CUD: cocaine use disorder. HC: healthy controls. INTs: intrinsic neural timescales. L: left; R: right. VN: visual network. SMN: somatomotor network. DAN: dorsal attention network. LN: limbic network. FPN: frontoparietal control network. DMN: default mode network. F-PNRC: missed trial rate under the congruent condition. F-PNRI: missed trial rate under the incongruent condition. * *p* < 0.05. ** *p* < 0.01. *** *p* < 0.001. ns: not significant.

### Cognitive consequences of distributed timescale dysregulation in CUD

To explore the potential relationship between inhibitory control and INTs, this study examined performance on the flanker and go/no-go tasks. Although no significant group differences were found in either task (Table S2), task performance in the CUD group showed significant association with INTs (Figure 3B). For the flanker task, INTs in the VN (*r* = 0.378, *p* = 0.034), SMN (*r* = 0.461, *p* = 0.007), DAN (*r* = 0.507, *p* = 0.002), and FPN (*r* = 0.445, *p* = 0.010) were positively correlated with the missed trial rate under congruent conditions. Under incongruent conditions, INTs in the SMN (*r* = 0.431, *p* = 0.013), DAN (*r* = 0.476, *p* = 0.005), and FPN (*r* = 0.456, *p* = 0.008), but not the VN (*r* = 0.319, *p* = 0.079), were positively correlated with the missed trial rate.

At the regional level, INTs in the left middle temporal gyrus (*r* = 0.355, *p* = 0.049) were positively correlated with the missed trial rate under incongruent conditions. Regarding error rate, INTs in the VN (congruent: *r* = 0.392, *p* = 0.028; incongruent: *r* = 0.436, *p* = 0.012) were positively correlated with error rate under both congruent and incongruent conditions, whereas INTs in the left fusiform gyrus (*r* = 0.393, *p* = 0.028) were positively correlated only under incongruent conditions. All reported *p*-values survived FDR correction. No significant correlations were found for the go/no-go task.

In HC, no significant correlations were observed (Figure S4). These findings suggest that in individuals with CUD, attentional functioning is closely linked to prolonged INT abnormalities. Specifically, higher INTs are associated with greater difficulties in inhibitory control.

### Mediating role of DAN INTs in predicting inhibitory control

To clarify how altered regional INTs predict inhibitory control, a mediation analysis was conducted to test whether INTs in large-scale networks (SMN, DAN, FPN) mediate the relationship between regional INT alterations and inhibitory control. As shown in Figure 3B, regression coefficients between left middle temporal gyrus and DAN INTs were statistically significant (a = 0.74, *p* < 0.001), whereas regression coefficients between the missed trial rate under incongruent condition and DAN INTs were not significant (b = 0.32, *p* = 0.122). The total effect and bootstrapped indirect effect were statistically significant (c = 0.49, *p* < 0.001; ab = 0.23, 95% CI [0.01, 0.51]), whereas the direct effect was not significant (c’ = 0.26, *p* = 0.204). These finding indicate that DAN INTs mediate the relationship between left middle temporal gyrus INTs and inhibitory control in CUD. In contrast, no mediation effects were observed when the SMN and FPN brain networks were included as mediators (Figure S5). Overall, these results suggest that prolonged INTs in DAN may represent a key neural mechanism underlying inhibitory control deficits in individuals with CUD.

### Altered INTs correlate positively with gamma-aminobutyric acid type A (GABAa) receptor and noradrenaline transporter (NAT) maps

To investigate the molecular underpinnings of INT alterations in CUD, t-values from the whole-brain analysis were extracted. INT differences (t-values) showed significant correlations with GABAa receptor (*r* = 0.43, *p* = 0.026, after FDR corrected) and NAT maps (*r* = 0.27, *p* = 0.026, after FDR corrected; Figure 4A). In analyses without GSR, the INT changes in CUD were also significantly correlated with GABAa receptor, NAT, and μ-opioid receptor (MU) distributions (Figure S6).

**Fig. 4.**
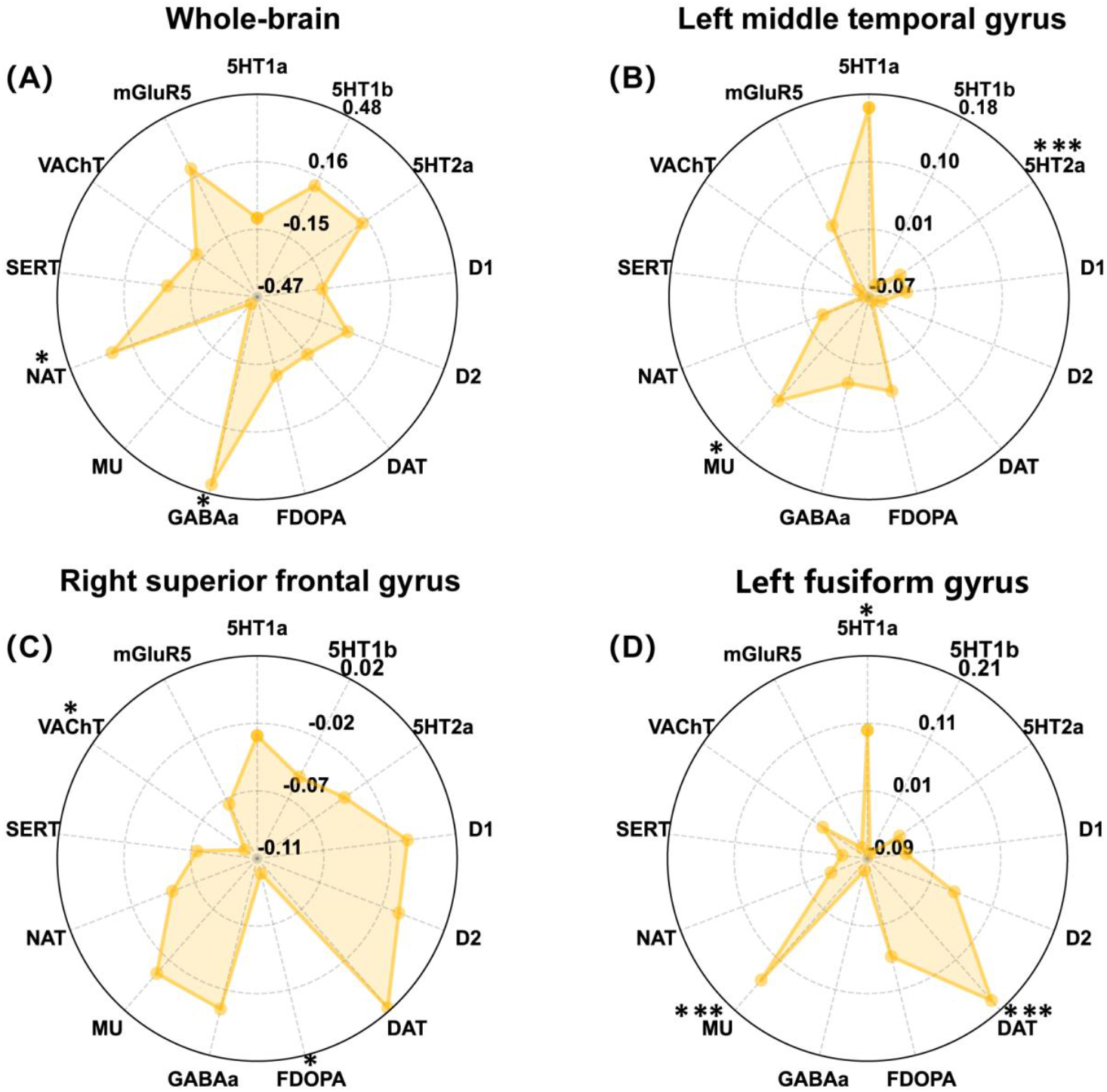
INTs specialization patterns associated with neurotransmitters. **(A)** Association between whole-brain INTs changes and neurotransmitters. **(B)** Association between INTs changes in the left middle temporal gyrus and neurotransmitters. **(C)** Association between INTs changes in the right superior parietal gyrus and neurotransmitters. **(D)** Association between INTs changes in the left fusiform gyrus and neurotransmitters. 5HT1a: serotonin 5-hydroxytryptamine receptor subtype 1a. 5HT1b: serotonin 5-hydroxytryptamine receptor subtype 1b. 5HT2a: serotonin 5-hydroxytryptamine receptor subtype 2a. D1: dopamine D1 receptor. D2: dopamine D2 receptor.DAT: dopamine transporter. FDOPA: fluorodopa. GABAa: gamma-aminobutyric acid type A receptor. MU: μ-opioid receptor. NAT: noradrenaline transporter. SERT: serotonin transporter. VAChT: vesicular acetylcholine transporter. mGluR5: metabotropic glutamate receptor 5. * *p* < 0.05, FDR corrected. ** *p* < 0.01, FDR corrected. *** *p* < 0.001, FDR corrected.

In the ROI analysis, INT differences in the left middle temporal gyrus were significantly correlated with 5-hydroxytryptamine 1A (5HT1a; *r* = 0.16, *p* < 0.001, after FDR corrected) and MU map (*r* = 0.10, *p* = 0.029, after FDR corrected; Figure 4B). INT difference in the right superior parietal gyrus was significantly correlated with fluorodopa (FDOPA) (*r* = -0.10, *p* = 0.042, after FDR corrected) and the vesicular acetylcholine transporter (VAChT) maps (*r* = -0.10, *p* = 0.041, after FDR corrected; Figure 4C). In the left fusiform gyrus, INT differences in significantly correlated with 5HT1a (*r* = 0.10, *p* = 0.042, after FDR corrected), dopamine transporters (DAT) (*r* = 0.19, *p* < 0.001, after FDR corrected), and MU (*r* = 0.15, *p* < 0.001, after FDR corrected; Figure 4D). Overall, these findings suggest that the molecular mechanisms underlying abnormal INT alterations in CUD are primarily associated with GABAa and NAT pathways.

### Assessment of classification performance using voxel features

To evaluate the potential of INTs as neuroimaging biomarkers for CUD, INTs from abnormal brain regions were extracted and used as input features for classification analysis. As shown in Table S3 and Figure 5, the support vector machine (SVM)(mean accuracy: 75.6%, *p*_permute_ < 0.001; mean AUC: 0.843, mean sensitivity: 79.9%, mean specificity: 72.7%) outperformed logistic regression (LR) (mean accuracy: 72.0%, *p*_permute_ = 0.008; mean area under the curve (AUC): 0.812, mean sensitivity: 78.6%, mean specificity: 65.6%) and k-nearest neighbor (KNN) (mean accuracy: 71.9%, *p*_permute_ = 0.007; AUC: 0.787, mean sensitivity: 86.5%, mean specificity: 58.7%) models, demonstrating the best classification performance.

**Fig. 5.**
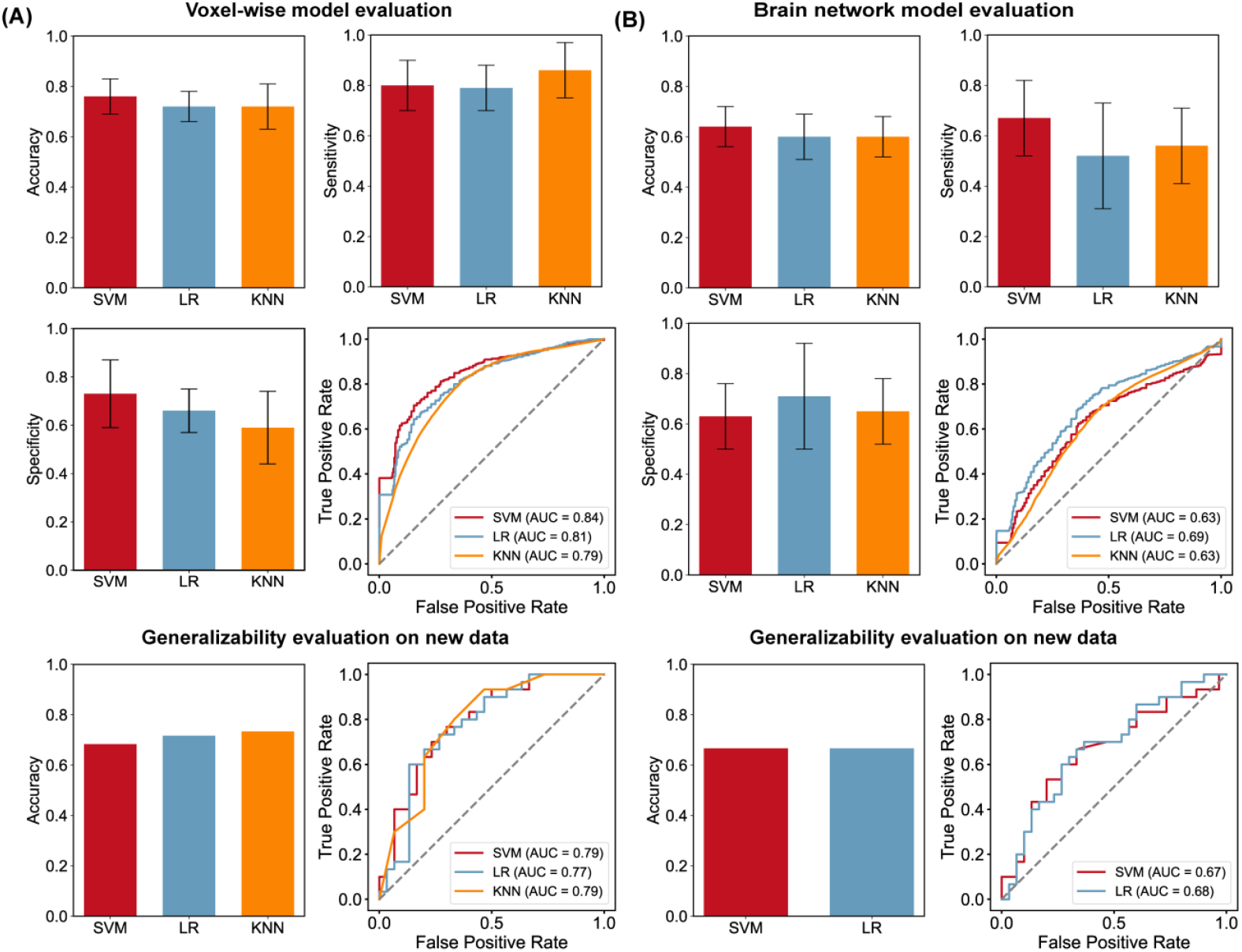
Classification of CUD and HC based on INTs features. **(A)** Evaluation of voxel-based feature classifiers. **(B)** Evaluation of brain network feature classifiers. SVM: support vector machine. LR: logistic regression. KNN: k-nearest neighbor. AUC: area under curve. Error bars represent standard deviation.

In analyses without GSR, however, the LR (mean accuracy: 71.2%, *p*_permute_ = 0.009; mean AUC: 0.754, mean sensitivity: 73.8%, mean specificity: 68.5%) achieved the best classification performance compared with SVM (mean accuracy: 69.7%, *p*_permute_ < 0.001; mean AUC: 0.756, mean sensitivity: 74.0%, mean specificity: 66.4%) and KNN (mean accuracy: 66.6%, *p*_permute_ = 0.044; mean AUC: 0.721, mean sensitivity: 67.8%, mean specificity: 66.1%) models, (Table S4 and Figure S7).

### Assessment of classification performance using brain network features

To further evaluate the ability of brain network features to differentiate HC from individuals with CUD, significantly altered network-level INTs were used as input features to construct SVM, LR, and KNN classification models. As shown in Figure 5 and Table S3, the SVM (mean accuracy: 64.2%, *p*_permute_ = 0.021; mean AUC: 0.629, mean sensitivity: 66.9%, mean specificity: 62.9%) demonstrated higher accuracy than LR (mean accuracy: 59.5%, *p*_permute_ = 0.015; mean AUC: 0.693, mean sensitivity: 52.2%, mean specificity: 71.2%), whereas the KNN model did not achieve statistical significance (mean accuracy: 60.1%, *p*_permute_ = 0.092; mean AUC: 0.629, mean sensitivity: 55.8%, mean specificity: 65.3%). In analyses without GSR, the classification performance of SVM (mean accuracy: 58.0%, *p*_permute_ = 0.115; mean AUC: 0.567, mean sensitivity: 55.7%, mean specificity: 63.2%), LR (mean accuracy: 54.4%, *p*_permute_ = 0.090; mean AUC: 0.664, mean sensitivity: 49.9%, mean specificity: 66.5%), and KNN (mean accuracy: 58.0%, *p*_permute_ = 0.163; mean AUC: 0.576, mean sensitivity: 47.4%, mean specificity: 69.4%) did not reach statistical significance (Table S4 and Figure S7).

### Evaluation of the generalizability of classification models

To assess the generalizability of the classification models, five classifiers were tested on an independent dataset to distinguish CUD from HC. For voxel feature classifiers, the SVM (accuracy: 68.3%, AUC: 0.790, sensitivity: 53.3%, specificity: 83.3%) demonstrated the best generalization performance compared with LR (accuracy: 71.7%, AUC: 0.770, sensitivity: 76.7%, specificity: 66.7%) and KNN (accuracy: 73.3%, AUC: 0.790, sensitivity: 93.3%, specificity: 53.3%) (Figure 5A). For brain network-based models, both SVM and LR achieved comparable classification accuracy of 66.7% with AUC values above the reported threshold 0.650. Overall, the machines learning results indicate that the SVM classifier based on voxel features achieved the highest performance among the tested models, highlighting the potential of INTs as neuroimaging biomarkers for CUD.

## Discussion

We used rs-fMRI data to investigate abnormal INTs changes in CUD and their associations with molecular expression. We also evaluated and validated the potential of INTs as diagnostic markers for CUD. INTs in the left pallidum and DMN correlated positively with abstinence duration, suggesting persistence of abnormalities during recovery. Altered INTs in the left middle temporal gyrus disrupted DAN dynamics, contributing to inhibitory control deficits in CUD. At the molecular level, these INT alterations were primarily associated with GABAa receptors and the NAT. Machine learning models using INTs successfully distinguished CUD from HC, with accuracy maintained above threshold in an independent dataset. These finding highlight neurodynamic alterations in CUD and their molecular underpinnings, providing novel insights into the pathophysiology of the disorder and supporting INTs as promising neuroimaging biomarkers.

### Prolonged INTs in CUD

In CUD, INTs were abnormally increased in the left middle temporal gyrus and left fusiform gyrus compared with HC. The left middle temporal gyrus, a key component of the DMN, supports semantic processing and exhibits higher betweenness centrality than in HC (*39*), reflecting its role in cross-regional integration under addiction. Both the middle temporal gyrus and DMN are critically involved in cue reactivity (*40*) and decision-making control (*41*–*43*).Under normal conditions, INTs maintain a balance between temporal segregation and integration (*17*). However, abnormally prolonged INTs in these regions may disrupt perceptual integration, impairing the accurate recognition and synthesis of perceptual information. Such disruption may contribute to altered semantic comprehension and emotional perception in CUD (*44, 45*).

INTs were also elevated in the right superior parietal gyrus, left pallidum, and left cerebellum_6 in CUD compared with HC. The superior parietal gyrus integrates spatial information and guides motor planning (*46*), whereas the pallidum–cerebellum circuit contributes to the refinement of motor execution (*47*–*49*). Prolonged INTs in these regions may disrupt the predictive processing of the brain by reducing the accuracy of internal predictions generated in sensory regions regarding motor intentions (*17, 50*). An abnormal increase in temporal scales could diminish predictive accuracy (*17, 51*), and amplify differences between intended and executed actions. These discrepancies may in turn impair the adjustment of subsequent motor programs in the superior parietal gyrus, ultimately leading to imprecise motor coordination (*52, 53*).

The role of the cerebellum in motor timing is well established (*54*), and timing deficits observed in substance use disorders may arise from such abnormalities in CUD (*55, 56*). Future studies could integrate multimodal neuroimaging with behavioral measures to clarify cerebellar contributions to time perception deficits in CUD. Elevated INTs, which reflect prolonged local neural activity and stronger autocorrelation (*17*), may stabilize connectivity within and between the DMN and SMN (*16*). However, this stability occurs at the expense of temporal flexibility (*57*), leading to rigid cross-network integration. Such rigidity can compromise the adaptive responses of the brain to external inputs.

The positive correlation between INTs and abstinence duration suggests persistent cocaine-induced neurodynamic disruptions, consistent with observation in other substance use disorders (*56, 58*–*60*). In CUD, abnormal INTs in the left middle temporal gyrus and related networks were associated with inhibitory control measures, indicating that prolonged INTs may contribute to impaired response inhibition (*57, 61*). Longer INTs are generally linked to broader attentional focus, whereas shorter INTs support more focused attention (*62*). In CUD, mediation analysis demonstrated that increased INTs in the left middle temporal gyrus altered DAN dynamics, which in turn predicted inhibitory control deficits. These regional disruptions may propagate through whole-brain connectivity, impairing spatiotemporal integration and contributing to cognitive dysfunction.

Correlation and mediation analyses together suggest that impaired attentional focus in CUD may manifest as increased INTs. Future studies should employ graph theory and directional connectivity approaches to validate the driving role of the middle temporal gyrus–DAN pathway.

Abnormal regions identified in CUD overlapped with those reported in male tobacco use disorder. However, the effects on INTs differed: INTs were increased in CUD but decreased in TUD (*28*). This pattern aligns reports of opposite blood-oxygen-level– dependent activity in methamphetamine use disorder and cocaine dependence (*31*). These findings suggest that addictive substances may induce substance-specific neuroplastic effects on brain networks by modulating INTs directionality of INT alterations. This study also systematically examined the impact of GSR. Although some left hemisphere findings varied in analyses without GSR, the core regions including the VN, SMN and DMN remained unaffected. GSR reduces motion- and respiration-related noise, thereby improving the signal-to-noise ratio and revealing neural activity that might otherwise be obscured (*63, 64*). Because GSR has been widely applied in rs-fMRI studies of INTs and can enhance their stability by improving signal quality, its use is supported in the present analysis (*28*–*30, 65*).

### Correlations with molecular features

INT alterations in CUD, relative to HC, were associated primarily with GABAa receptors and the NAT. Cocaine neurotoxicity involves abnormal elevation of synaptic dopamine, serotonin, and norepinephrine, leading to neurological damage (*66*). GABAa receptors modulate compulsive cocaine-seeking behavior and may therefore contribute to the observed INT abnormalities (*67*). In socio-emotional regions (*68, 69*), INTs changes in the left middle temporal gyrus and fusiform gyrus were associated with 5HT1a and MU receptors. INT alterations in the right superior parietal gyrus and left fusiform gyrus were linked to the dopaminergic system, including the DAT and FDOPA.

Addiction involves multiple neurotransmitter systems dopamine, glutamate, serotonin, and GABAa working in concert (*70*). Among these, serotonin and GABAa systems act as inhibitory modulators of emotional regulation (*71*). Notably, MU receptors may enhance dopamine release by inhibiting GABAa interneurons, suggesting that with dopamine–MU synergy could shape the temporal dynamics of local neural activity(*72*).

INT changes in the superior parietal gyrus were also associated with the VAChT, key component of cholinergic neurotransmission. VAChT deficiency alters dendritic morphology and electrophysiological properties, thereby affecting temporal integration (*19, 73*). These findings align with prior reports on tobacco use disorder (*28*), emphasizing the role of acetylcholine in addiction. By regulating neurotransmitter release and dendritic properties, the vesicular acetylcholine transporter may influence temporal integration at the network level. Future studies should investigate causal links between neurotransmitter release, dendritic plasticity, and INT alterations in addiction, which could clarify how molecular mechanisms shape large-scale neural dynamics in substance use disorders.

### INTs as a potential biomarker for CUD

This study applied supervised machine learning methods to evaluate the diagnostic potential of INT features. Classifiers based on voxel-wise INTs effectively distinguished CUD from HC. SVM and LR model based on voxel-level INT features achieved classification accuracies of 75.5% and 72.0%, both exceeding the widely accepted threshold for good classifiers in clinical research (AUC > 0.800) (*74*). Using brain network-level INT features, SVM and LR models achieved classification accuracies of 64.2% and 59.5%, respectively. In analyses without GSR, both voxel-based SVM and LR models-maintained accuracies exceeding 70.0%. However, classification performance using brain network features did not reach statistical significance, likely owing to the limited number of available features in the non-GSR data (only two network features). This finding highlights the superior discriminative power of voxel-level INT, particularly when extracted after GSR.

Classifier generalization was validated in an independent dataset, with accuracy rates consistently exceeding 65.0%, providing additional insights beyond previous studies (*75*). Voxel-based features achieved higher classification accuracy than brain network features (75.5% vs 64.2%), suggesting that localized neural dynamics may serve as more sensitive biomarkers. Robustness was further confirmed across diverse algorithms and datasets, indicating that INT based classification is not dependent on a specific algorithm.

Compared with structural imaging features (AUC = 0.8) (*76*), INTs demonstrated comparable potential as biomarkers for CUD. These findings extend the application of INTs to substance addiction and highlight their promise for personalized CUD diagnosis, advancing the development of objective neuroimaging tools.

The present study has several limitations. First, the relationship between INTs and neurotransmitters was inferred indirectly. To achieve a more comprehensive understanding of the relationship between brain alterations relate to molecular biology in CUD, future investigations integrating fMRI with PET data are needed. Second, the abstinence duration of participants in this study was relatively short. Future research should include individuals with wider range of abstinence periods, ideally categorized into distinct phases, to clarify the temporal trajectory of INT alterations during recovery.

Despite these limitations, this study systematically investigated INTs abnormalities and their associated molecular features in CUD. Abnormalities were primarily concentrated in the VN, SMN, and DMN. Elevated INTs in specific regions were linked to disrupted DAN dynamics and impaired inhibitory control in CUD. Supervised machine learning further validated INTs as potential diagnostic biomarkers for CUD. These findings advance our understanding of the pathophysiology of CUD and support the development of objective neuroimaging biomarkers and potential therapeutic targets.

## Materials and Methods

### Experimental procedures

For the resting-state fMRI, the participants needed to stay awake and keep their eyes open, watching a cross at the center of the screen. If they closed their eyes for more than ten seconds, they were reminded to stay awake, and the process was restarted. Behavioral task analysis is important as it reveals cognitive processing features related to CUD and offers insights into cognitive control. Combining rs-fMRI with behavioral tasks helps study the relationship between resting-state brain function and cognitive processing in CUD patients. After the MRI scan, each participant completed a 45-minute cognitive test, which included several tasks such as Berg’s card sorting task, flanker task, go/no-go task, letter-number sequencing, digit span backward, Iowa gambling task, and tower of London. For more detailed information on the questionnaires and experimental procedures, please see the previous study (*77*).

### Participants

The data analyzed in this study were obtained from the SUDMEX CONN dataset (*77*). This dataset comprised 138 participants, including 64 HC and 74 CUD participants. According to the Declaration of Helsinki, all participants provided informed consent both in writing and verbally. In this dataset, the diagnosis of CUD participants was performed using the MINI International Neuropsychiatric Interview - Plus Spanish version 5.0.0. The CUD patients had used cocaine for at least one year, with a frequency of at least three days per week with no more than 60 continued days of abstinence in the past year.

Additional inclusion and exclusion criteria can be found in the description of the published dataset (*77*). In the current study, a total of 24 participants (CUD = 16, HC = 8) were excluded due to missing functional imaging data (n = 3), incomplete basic demographic information (n = 5), unavailable cocaine uses information (n = 3), a history of substance uses in the HC group (n = 3), or excessive head motion (n = 10). Referring to previous studies (*16, 78*), this study matched participants based on age, education, and gender by excluding older, less educated participants from the CUD group and younger, more educated participants from the HC group. A total of 90 participants were included in the following analyses. Based on the 2.5 standard deviation criterion (*79*), two participants were shown as having outlier whole-brain INTs values. Consequently, a total of 88 participants were included in the final statistical analyses (CUD group = 44, HC group = 44). The Liaoning Normal University Human Research Institutional Review Board approved the secondary data analysis.

### MRI data acquisition and preprocessing

Imaging data were collected on a Philips Ingenia 3T scanner (Philips Healthcare, Best, The Netherlands) with a 32-channel dS Head coil. The rs-fMRI was obtained using a gradient recalled echo planar imaging (EPI) sequence with the following parameters: repetition time = 2000 ms, echo time = 30.001 ms, flip angle = 75°, matrix = 80 × 80, field of view = 240 mm^2^, voxel size = 3 × 3 × 3 mm. The slices were acquired in an interleaved (ascending) order, number of slices = 36, phase encoding direction = anterior-posterior.

The rs-fMRI session lasted for 10 minutes, during which 300 volumes were collected. The rs-fMRI data were preprocessed using the Data Processing and Analysis of Brain Imaging (DPABI) Toolbox (*80*), which operates on the MATLAB platform. The preprocessing steps mainly included the following: first, the removal of the first 10 volumes, followed by slice timing correction and head motion correction. Then, the fMRI images were spatially normalized to the standard EPI template, resampled to a voxel size of 3 × 3 × 3 mm^3^. After normalization, nuisance covariate regression was performed to remove the effects of Friston-24 motion parameters, white matter signals, cerebrospinal fluid signals, and global signal. Spatial smoothing was applied using a Gaussian kernel with a full-width at half-maximum of 6 mm. Finally, a band-pass filter between 0.01 and 0.08 Hz was applied to remove noise from both low and high frequencies. Participants were excluded if their head motion exceeded a maximum displacement of 3 mm or a rotation of 3°. Additionally, this study calculated the mean framewise displacement (FD) of each participant. There was no significant difference of mean FD between CUD and HC (*t*_(88)_ ≤ 0.001, *p* = 0.99, Cohen’s d ≤ 0.001).

### Calculation of intrinsic neural timescales

According to previous studies (*25, 81*), the size of the INTs was estimated by calculating the area under the curve of the autocorrelation function (ACF). ACF was used to measure the temporal correlation of neural activity in local brain regions. A higher ACF indicated greater stability in the neural activity patterns of that region. The fMRI signal from each voxel was extracted, and the ACF was estimated using the following formula:

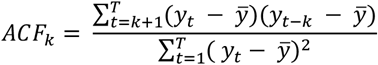

where the rs-fMRI signal is represented by y, and 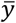is the mean signal across timepoints. T indicates the number of time points, and k is the time lag. Subsequently, the sum of the ACF values during the initial period when the ACF remained positive was calculated. The initial positive period included all time points from the initial lag until the ACF first reached zero. This whole-brain map was used as the INTs distribution map, with each voxel storing the INTs value of the corresponding brain region. In addition, this study calculated the functional brain network INTs based on whole-brain INTs maps.

According to Yeo’s 7-network standard atlas (MNI152 space), the seven brain networks were defined as follows: the VN, SMN, DAN, VAN, LN, FPN, and DMN (*82*). To estimate the INTs for each brain network, based on previous studies (*81, 83*), we calculated the average INTs of all voxels within each of the seven networks. Whole-brain INTs and brain network INTs were visualized using BrainNet Viewer (*84*) and the R package ggseg (*85*), respectively. The code for extracting INTs is from GitHub (https://github.com/takamitsuwatanabe/IntrinsicNeuralTimescale) (*81*).

### Spatial correlations between altered INT and neurotransmitters

To investigate the potential physiological mechanisms underlying cocaine-induced abnormal INTs, the Juspace toolbox was used to evaluate whether INTs changes in CUD relate to specific neurotransmitter receptors/transporters (*86*). 13 types of neurotransmitter receptors/transporters were selected from the JuSpace toolbox. These include the 5-hydroxytryptamine receptor subtypes 1a, 1b, and 2a (5HT1a, 5HT1b, and 5HT2a) (*87, 88*), dopamine D1 and D2 receptors (D1 and D2) (*89, 90*), DAT (*91*), FDOPA (*92*), GABAa (*93*), (MU (*90*), NAT (*94*), serotonin transporter (SERT) (*88*), VAChT (*90*), and metabotropic glutamate receptor 5 (mGluR5) (*90*). The neurotransmitter receptors and transporters selected were the most commonly used in previous research, with details provided in Table S5. The INTs differences (t-values) between CUD and HC were used as inputs for Spearman correlation analyses with PET/SPECT imaging (*95*). This study employed the Neuromorphometrics atlas, which serves as the default anatomical atlas, to extract mean regional values from the input images for correlation with the regional values of neurotransmitter maps. The exact *p*-values were determined using 1,000 permutations, with spatial autocorrelation adjustments. To correct for multiple comparisons of the exact *p*-values related to the 13 types of neurotransmitters, the FDR was applied, with a significance threshold set at *p* < 0.05. Furthermore, this study also extracted neurotransmitter densities from brain regions showing significant differences in INTs between CUD and HC. Spearman correlation analyses were then conducted to examine the relationship between neurotransmitter densities and changes in INTs within these regions. This allowed us to explore whether changes in INTs within key regions are associated with neurotransmitter densities.

### Classification analysis using machine learning

To evaluate the discriminatory power of INTs between CUD and HC, this study constructed a radial basis function kernel SVM model using the Python package scikit-learn. Two SVM models were evaluated: (1) voxel-based features; (2) brain network features. First, voxel-wise INTs with significant differences (after TFCE-FDR correction) or brain network INTs with significant differences (after FDR correction) were selected as input features, respectively. The total number of voxel-based features reached 2,389, which far exceeded the sample size. It is necessary to perform dimensionality reduction to avoid the curse-of-dimensionality or the small-n-large-p problem (*96*). We standardized all feature data using a Min-Max scaler and applied principal component analysis to the voxel-based features, retaining 55 features that accounted for 90% of the variance. Then, this study employed five-fold cross-validation with 100 iterations and used grid search in each iteration to find the optimal parameters. Finally, we assessed statistical significance using 10,000 permutations, with a threshold of *p* < 0.05. The classification performance of INTs was evaluated by reporting the AUC of the receiver-operating characteristic curves, accuracy, sensitivity, and specificity. To better evaluate the stability of INTs’ classification performance and examine that the classification efficacy of INTs does not rely on a single classifier, this study also constructed LR and KNN classifiers.

### Statistical analysis

In this study, independent sample t-test and chi-square test were employed to analyze differences in demographic information and cocaine use data between the CUD and HC groups. For the INTs metric, independent sample t-tests were performed on the INTs maps of the two groups using the DPABI toolbox, with age, gender, and education as covariates. The statistically significant threshold was set at *p* < 0.05 (two-tailed), following TFCE-FDR correction with 1,000 permutations (*97*).

Two-sample t-tests were also conducted for each brain network to identify those showing statistically significant differences in INTs between the CUD and HC groups, with a significance threshold of *p* < 0.05. The FDR correction was applied to control for multiple comparisons. To assess the overlap between significant regions and the seven brain networks across the two groups, the DC was computed for the cluster and its corresponding brain network (*98*). The DC is a metric used to quantify the degree of overlap between two regions. The average INTs for each ROI was obtained using the ROI signal extraction function implemented in DPABI.

Spearman correlation analysis was performed to assess the associations between INTs (including ROI INTs and brain network INTs) and clinical variables in CUD. In addition, previous studies have proposed that individuals with substance use disorders exhibit abnormal neural activity in the parietal lobe during inhibitory control tasks (*99*). This aberrant neural activity may be associated with susceptibility to substance use (*9*). Therefore, the current study analyzed behavioral performance in the flanker task and the go/no-go task. Detailed procedures for these tasks can be found in prior research (*77*). FDR correction was applied to the correlation analysis results.

This study employed mediation analysis to investigate the relationships among ROI INTs, brain network INTs, and clinical information, using the PROCESS v4.2 (*100*) in SPSS 26 software. ROI INTs were set as the independent variable, brain network INTs served as the mediating variable, and clinical information was treated as the dependent variable. To evaluate indirect effects, the bootstrapping method with 5000 iterations was employed. If the 95% confidence interval (CI) did not include zero, the mediation effect was regarded as significant (*p* < 0.05). To ensure that the observed mediation effect reflects a true intermediate mechanism rather than shared variance due to overlapping regions, this study excluded the INTs of voxels overlapping between the ROIs and the brain networks, and then constructed the mediation model.

### Reproducibility analysis

Two methodological strategies were employed to ensure the robustness of the primary findings and the stability of machine learning classification. First, the analysis was conducted on data without GSR, and to further assess the impact of GSR, covariates such as age, gender, and education were excluded from the analysis. The results were comprehensively documented in the Supplementary Materials. Second, this study also selected CUD participants from the pre-test data of the SUDMEX TMS dataset (*101*) and constructed a new dataset with 30 CUD and 30 HC participants, matched on age, education, and gender. Demographic information can be found in the Supplementary Materials (Table S6). The constructed classifiers were then applied to the new dataset to evaluate their generalizability.

## Acknowledgments

We thank Eduardo A. Garza-Villarreal and colleagues for publicly sharing the SUDMEX TMS and SUDMEX CONN datasets. Their valuable efforts in data collection and open access have been instrumental to the success of this study.

## Author contributions

Conceptualization: X.T. and M.Z.

Methodology: X.T., T.D., J.S., M.Z., and W.L.

Investigation: X.T., T.D., J.S., and M.Z.

Visualization: X.T. and M.Z.

Supervision: M.Z. and W.L.

Funding acquisition: M.Z. and W.L.

Writing—original draft: X.T. and M.Z.

Writing—review & editing: X.T., T.D., J.S., Y.S., W.L., and M.Z.

## Data and materials availability

The datasets used in this study are publicly available. The SUDMEX CONN datasets can be accessed at https://openneuro.org/datasets/ds003346. The corresponding clinical and cognitive data are available on Zenodo (https://doi.org/10.5281/zenodo.5123331). The SUDMEX TMS dataset can be also accessed at https://openneuro.org/datasets/ds003037/versions/2.1.0. The corresponding clinical and cognitive data are available on Zenodo (https://doi.org/10.5281/zenodo.10409461).

