## Supplementary Materials for "Abnormally increased intrinsic neural timescales in sensory and default mode networks in cocaine use disorder"

Xiaoling Tang et al.

Dr. Wenbo Luo,

**This PDF file includes:**

Figs. S1 to S7
Tables S1 to S6

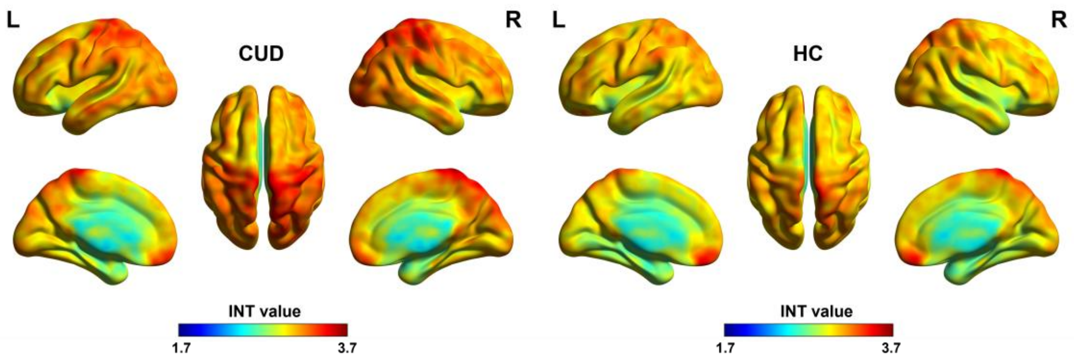


**Fig. S1. The distribution of INTs and differences between CUD and HC groups in the data without global signal regression.** CUD: cocaine use disorder. HC: healthy controls. INTs: intrinsic neural timescales.

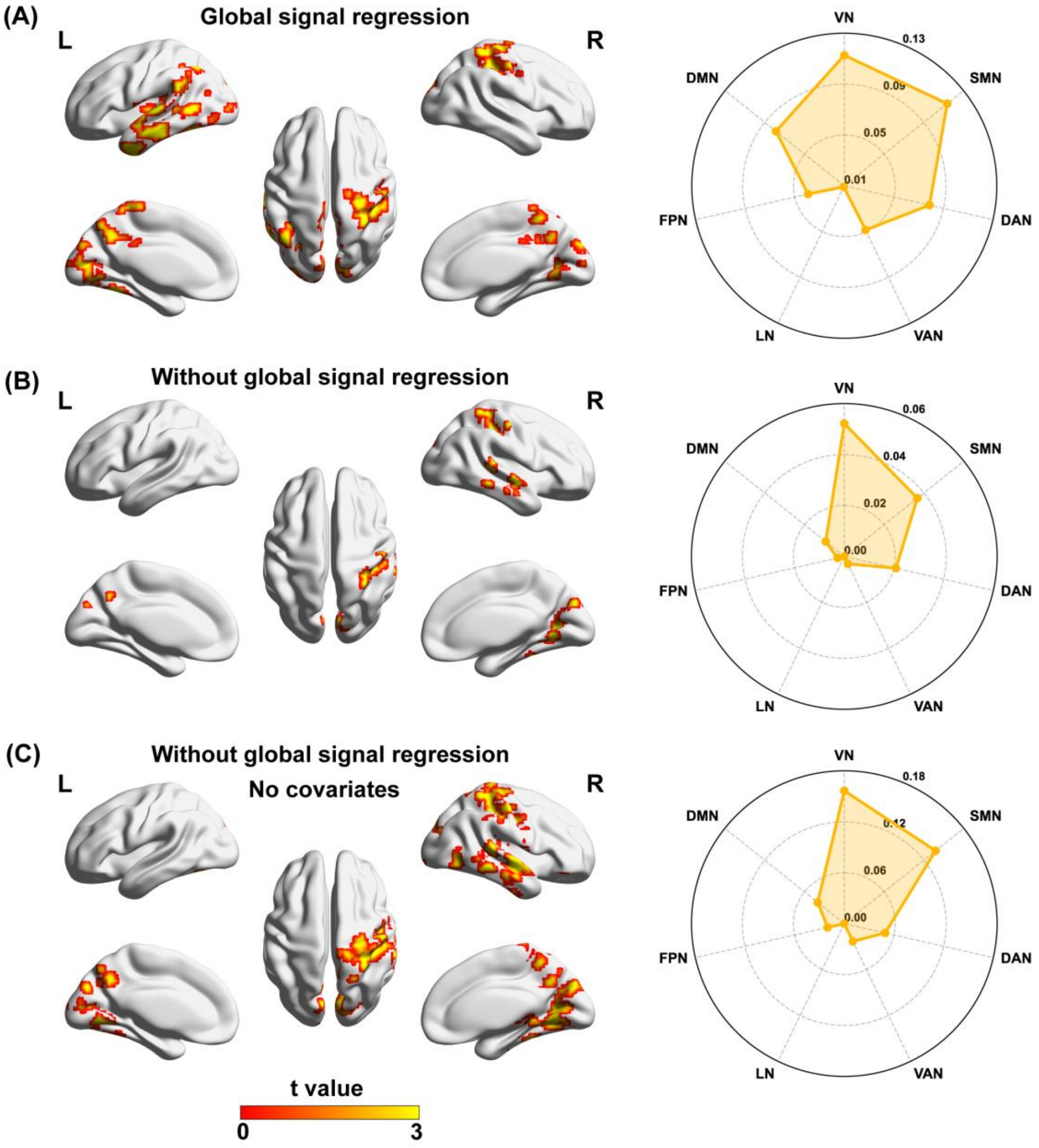


**Fig. S2. The brain regions showing significant differences between whole-brain analyses with and without global signal regression. (A)** The results of the whole-brain analysis with global signal regression. **(B)** The results of the whole-brain analyis without global signal regression. **(C)** The results of the whole-brain analysis without global signal regression and without the inclusion of covariates.

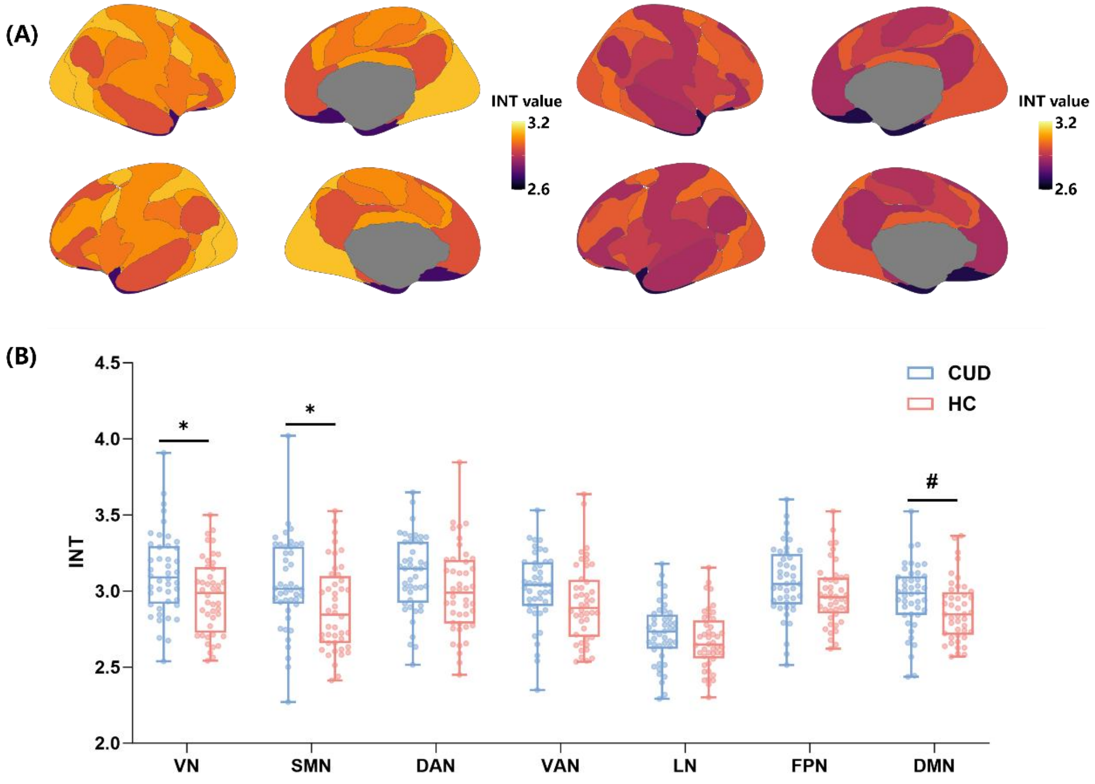


**Fig. S3. The distribution of INT and the differences between the CUD and HC groups in the data without global signal regression. (A)** The distribution pattern of the brain network INTs in the CUD and HC groups without global signal regression. **(B)** The averaged values of the brain network INTs in the CUD and HC groups without global signal regression. CUD: cocaine use disorder. HC: healthy controls. INTs: intrinsic neural timescales. VN: visual network. SMN: somatomotor network. DAN: dorsal attention network. VAN: ventral attention network. LN: limbic network. FPN: frontoparietal control network. DMN: default mode network. * $p < 0.05$, FDR corrected. # $p < 0.05$, uncorrected.

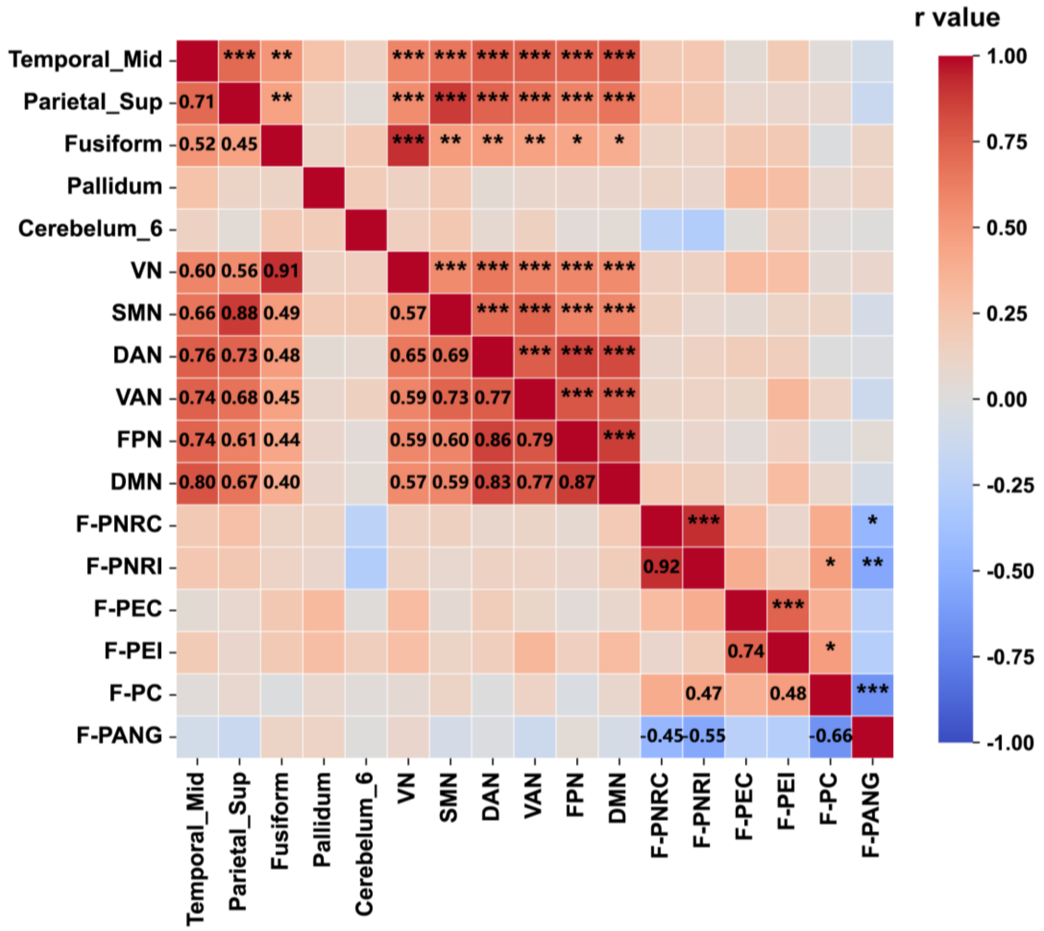


**Fig. S4. The relationship between INTs and behavioral information in HC.** HC: healthy controls. INTs: intrinsic neural timescales. F-PNRC: The rate of missed trials under congruent conditions. F-PNRI: The rate of missed trials under incongruent conditions. F-PEC: The error rate under congruent conditions. F-PEI: The error rate under incongruent conditions. F-PC: The error rate in No-Go trials. F-PANG: The accuracy rate in No-Go trials. VN: visual network. SMN: somatomotor network. DAN: dorsal attention network. VAN: ventral attention network. LN: limbic network. FPN: frontoparietal control network. DMN: default mode network. * $p < 0.05$. ** $p < 0.01$. *** $p < 0.001$.

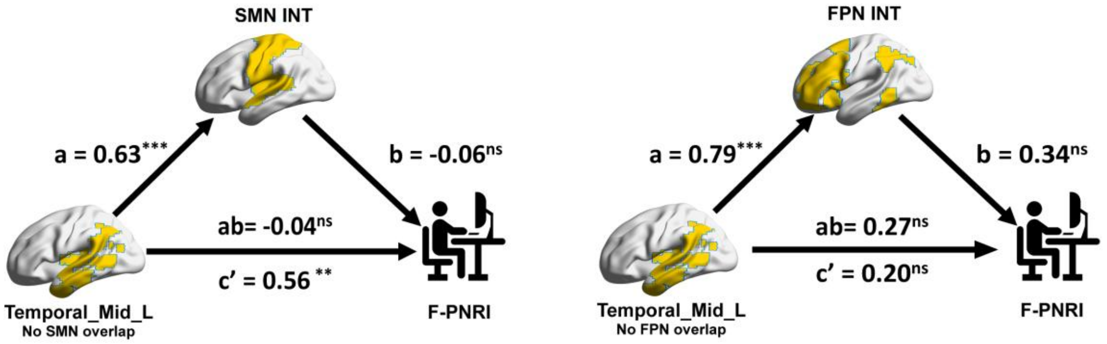


**Fig. S5. A mediation model with regional INTs as the independent variable, network INTs as the mediator, and behavioral information as the dependent variable.** INTs: intrinsic neural timescales. L: left; R: right. DAN: dorsal attention network. LN: limbic network. FPN: frontoparietal control network. F-PNRI: missed trial rate under the incongruent condition. * $p < 0.05$. ** $p < 0.01$. *** $p < 0.001$.

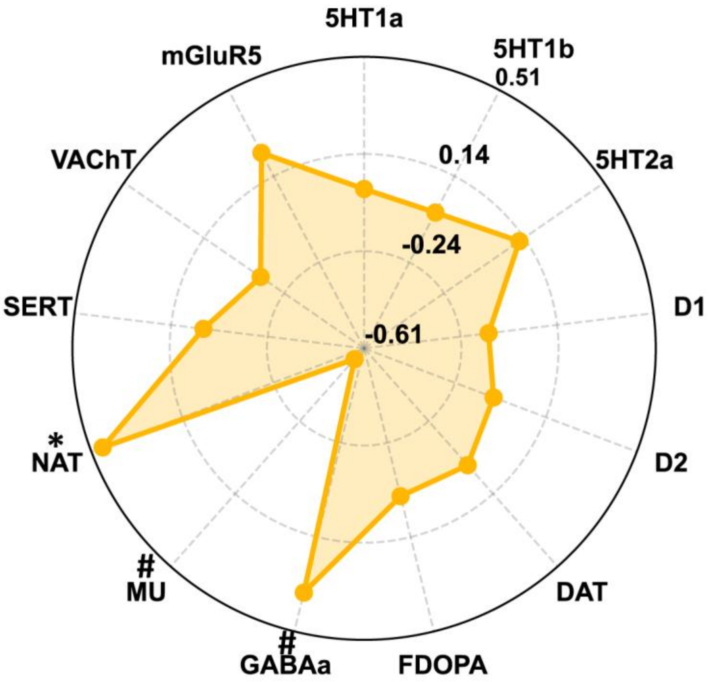


**Fig. S6. INT specialization patterns associated with neurotransmitters in the data without global signal regression.** 5HT1a: serotonin 5-hydroxytryptamine receptor subtype 1a. 5HT1b: serotonin 5-hydroxytryptamine receptor subtype 1b. 5HT2a: serotonin 5-hydroxytryptamine receptor subtype 2a. D1: dopamine D1 receptor. D2: dopamine D2 receptor. DAT: dopamine transporter. FDOPA: fluorodopa. GABAa: gamma-aminobutyric acid type A receptor. MU: μ-opioid receptor. NAT: noradrenaline transporter. SERT: serotonin transporter. VAChT: vesicular acetylcholine transporter. mGluR5: metabotropic glutamate receptor 5. * $p < 0.05$, FDR corrected. ** $p < 0.01$, FDR corrected. *** $p < 0.001$, FDR corrected. # $p < 0.05$, uncorrected.

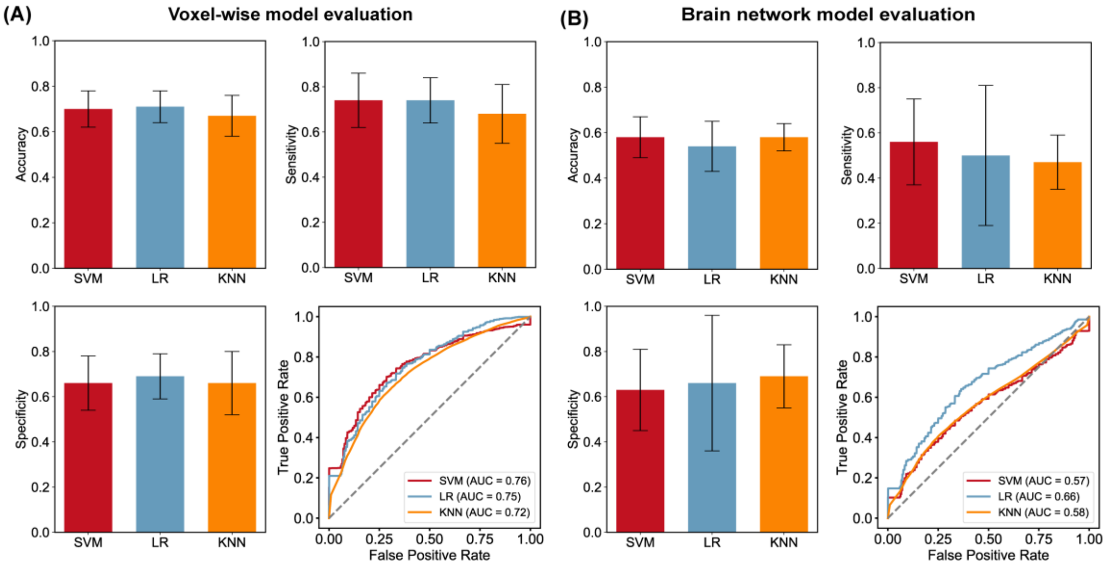


**Fig. S7. Classification of CUD and HC based on INTs features in data without global signal regression. (A)** Evaluation of voxel-based feature classifiers. **(B)** Evaluation of brain network feature classifiers. SVM: support vector machine. LR: logistic regression. KNN: k-nearest neighbor. AUC: area under curve. Error bars represent standard deviation.

**Table S1. Significant group differences in INTs in data without global signal regression.** AAL: Anatomical Automatic Labeling. MNI: Montreal Neurological Institute. CUD: cocaine use disorder. HC: healthy controls. INTs: intrinsic neural timescales. L: left; R: right.

| Region | AAL | Peak MNI Coordinate | | | Voxels | t value |
|---|---|---|---|---|---|---|
| | | X | Y | Z | | |
| **Postcentral_R** | 58 | 33 | -30 | 42 | 1033 | 4.34 |
| **Fusiform_R** | 56 | 33 | -39 | -18 | 1007 | 5.04 |
| **Cerebelum_6_L** | 99 | -24 | -54 | -30 | 91 | 3.49 |
| **Temporal_Sup_R** | 82 | 57 | -33 | 9 | 62 | 4.26 |
| **Thalamus_R** | 78 | 15 | -21 | 0 | 39 | 3.80 |

**Table S2. Statistical comparisons of behavioral tasks between the CUD and HC groups.** CUD: cocaine use disorder. HC: healthy controls. INTs: intrinsic neural timescales. F-PNRC: The rate of missed trials under congruent conditions. F-PNRI: The rate of missed trials under incongruent conditions. F-PEC: The error rate under congruent conditions. F-PEI: The error rate under incongruent conditions. F-PC: The error rate in No-Go trials. F-PANG: The accuracy rate in No-Go trials.

| **Variable** | **CUD** | **HC** | **t** | ***p*** | **Cohen's d** |
|---|---|---|---|---|---|
| **F-PNRC** | 0.09 ± 0.11 | 0.10 ± 0.04 | -0.63 | 0.53 | -0.15 |
| **F-PNRI** | 0.09 ± 0.12 | 0.12 ± 0.15 | -0.72 | 0.48 | -0.17 |
| **F-PEC** | 0.07 ± 0.12 | 0.05 ± 0.06 | 0.74 | 0.47 | 0.17 |
| **F-PEI** | 0.09 ± 0.12 | 0.08 ± 0.08 | 0.54 | 0.59 | 0.13 |
| **F-PC** | 0.07 ± 0.14 | 0.05 ± 0.07 | 0.56 | 0.58 | 0.13 |
| **F-PANG** | 0.93 ± 0.14 | 0.89 ± 0.23 | 1.01 | 0.31 | 0.24 |

**Table S3. The classification performance (M ± SD) of the six models.** SVM: support vector machine. LR: logistic regression. KNN: k-nearest neighbor. AUC: area under curve.

| **Model** | | **Train accuracy** | **Test accuracy** | **Test sensitivity** | **Test specificity** | **Test AUC** | ***p*** |
|---|---|---|---|---|---|---|---|
| **Voxel Feature** | **SVM** | 0.84 ± 0.03 | 0.76 ± 0.07 | 0.80 ± 0.10 | 0.73 ± 0.14 | 0.84 ± 0.06 | < 0.001 |
| | **LR** | 0.85 ± 0.02 | 0.72 ± 0.06 | 0.79 ± 0.09 | 0.66 ± 0.09 | 0.81 ± 0.06 | 0.008 |
| | **KNN** | 0.79 ± 0.05 | 0.72 ± 0.09 | 0.86 ± 0.11 | 0.59 ± 0.15 | 0.79 ± 0.08 | 0.007 |
| **Brainnet Feature** | **SVM** | 0.69 ± 0.04 | 0.64 ± 0.08 | 0.67 ± 0.15 | 0.63 ± 0.13 | 0.63 ± 0.16 | 0.021 |
| | **LR** | 0.64 ± 0.04 | 0.60 ± 0.09 | 0.52 ± 0.21 | 0.71 ± 0.21 | 0.69 ± 0.09 | 0.015 |
| | **KNN** | 0.68 ± 0.04 | 0.60 ± 0.08 | 0.56 ± 0.15 | 0.65 ± 0.13 | 0.63 ± 0.09 | 0.092 |

**Table S4. The classification performance (M ± SD) of the six models, constructed without global signal regression.** SVM: support vector machine. LR: logistic regression. KNN: k-nearest neighbor. AUC: area under curve.

| **Model** | | **Train accuracy** | **Test accuracy** | **Test sensitivity** | **Test specificity** | **Test AUC** | ***p*** |
|---|---|---|---|---|---|---|---|
| | **SVM** | 0.74 ± 0.03 | 0.70 ± 0.08 | 0.74 ± 0.12 | 0.66 ± 0.12 | 0.76 ± 0.12 | < 0.001 |
| **Voxel** | **LR** | 0.75 ± 0.02 | 0.71 ± 0.07 | 0.74 ± 0.10 | 0.69 ± 0.10 | 0.75 ± 0.08 | 0.009 |
| | **KNN** | 0.77 ± 0.05 | 0.67± 0.07 | 0.68 ± 0.13 | 0.66 ± 0.14 | 0.72 ± 0.08 | 0.044 |
| | **SVM** | 0.65 ± 0.05 | 0.58 ± 0.09 | 0.56 ± 0.19 | 0.63 ± 0.18 | 0.57 ± 0.15 | 0.115 |
| **Brainnet** | **LR** | 0.60 ± 0.04 | 0.54 ± 0.11 | 0.50 ± 0.31 | 0.66 ± 0.30 | 0.66 ± 0.08 | 0.090 |
| | **KNN** | 0.68 ± 0.05 | 0.58 ± 0.06 | 0.47 ± 0.12 | 0.69 ± 0.14 | 0.58 ± 0.08 | 0.163 |

**Table S5. Neurotransmitter density maps considered in the present study.**

| Receptor/ transporter | Neurotransmitter | Map |
|---|---|---|
| 5HT1a | Serotonin | 5HT1a_WAY_HC36 |
| 5HT1b | Serotonin | 5HT1b_az_hc36_beliveau |
| 5HT2a | Serotonin | 5HT2a_cimbi_hc29_beliveau |
| D1 | Dopamine | D1_SCH23390_c11 |
| D2 | Dopamine | D2_fallypride_hc49_jaworska |
| DAT | Dopamine | DAT_DATSPECT |
| FDOPA | Dopamine | FDOPA_f18 |
| GABAa | GABA | GABAa_flumazenil_hc16_norgaard |
| MU | Opioids | MU_carfentanil_hc39_turtonen |
| NAT | Noradrenaline | NAT_MRB_c11 |
| SERT | Serotonin | SERT_dasb_hc100_beliveau |
| VAChT | Acetylcholine | VAChT_feobv_hc18_aghourian |
| mGluR5 | Glutamate | mGluR5_abp_hc73_smart |

**Table S6. Demographic and clinical characteristics of both groups in the new data.** CUD: cocaine use disorder. CUD: cocaine use disorder. HC: healthy controls. FD: framewise displacements. CCQN: cocaine craving questionnaire now. NA: not applicable.

| Variable | CUD | HC | t or $\chi^2$ | *p* | Cohen's d or Phi |
|---|---|---|---|---|---|
| **Age (years)** | 35.23 ± 7.76 | 33.77 ± 7.61 | 0.74 | 0.46 | 0.19 |
| **Gender (male/female)** | 26/4 | 22/8 | 1.67 | 0.20 | 0.17 |
| **Education (years)** | 12.57 ± 2.76 | 12.43 ± 3.42 | 0.17 | 0.86 | 0.04 |
| **Mean FD (mm)** | 0.11 ± 0.05 | 0.12 ± 0.04 | -1.34 | 0.19 | -0.35 |
| **Duration of cocaine use (years)** | 10.47 ± 6.82 | NA | NA | NA | NA |
| **Age of first cocaine use (years)** | 23.77 ± 6.53 | NA | NA | NA | NA |
| **CCQN** | 144.20 ± 54.43 | NA | NA | NA | NA |
